# AxoMetric: A Rapid and Unbiased Tool for Automated Quantification of Axon Regeneration in Tissue Sections

**DOI:** 10.1101/2025.07.02.662816

**Authors:** Matthew C. Finneran, Tara Rhamani, Ismaël Valentin Salioski, Ligia B. Schmitd, Ryan Passino, Craig N. Johnson, Roman J. Giger

**Affiliations:** Neuroscience Graduate Program, University of Michigan Medical School, Ann Arbor, MI, USA; Cell and Developmental Biology, University of Michigan Medical School, Ann Arbor, MI, USA; Department of Neurology, University of Michigan Medical School, Ann Arbor, MI, USA; Faculty of Life Sciences, École Polytechnique Fédérale de Lausanne, Lausanne, CH

## Abstract

Recent advances in experimental strategies that promote axon regeneration in adult mammals lay the foundation for future therapies. Reliable and unbiased quantification of regenerated axons is challenging, yet essential for comparing the efficacy of individual treatments and identification of most efficacious combinatorial therapies. Here, we introduce *AxoMetric*, a user-friendly and freely available software for the rapid quantification of regenerated axons in longitudinal nerve tissue sections. *AxoMetric* automatically identifies and traces regenerated axons, generating quantitative measurements that closely match conventional manual quantification but with significantly greater speed. Key features include length-dependent axon quantification at defined intervals from the injury site and normalization of axon density to nerve diameter to account for anatomical variability. To facilitate high-throughput analysis, the software includes an image queuing function. Additional features of *AxoMetric* allow quantification of a range of labeled cellular structures. As a proof of concept, we demonstrate accurate quantification of regenerated axons in the optic nerve, retinal ganglion cells density in retinal flat-mounts, and regenerated axon bundles in injured sciatic nerves. Collectively, we introduce a new platform that is expected to streamline and standardize regenerative outcome assessments across diverse experimental conditions and laboratories.

**Graphical Abstract:** 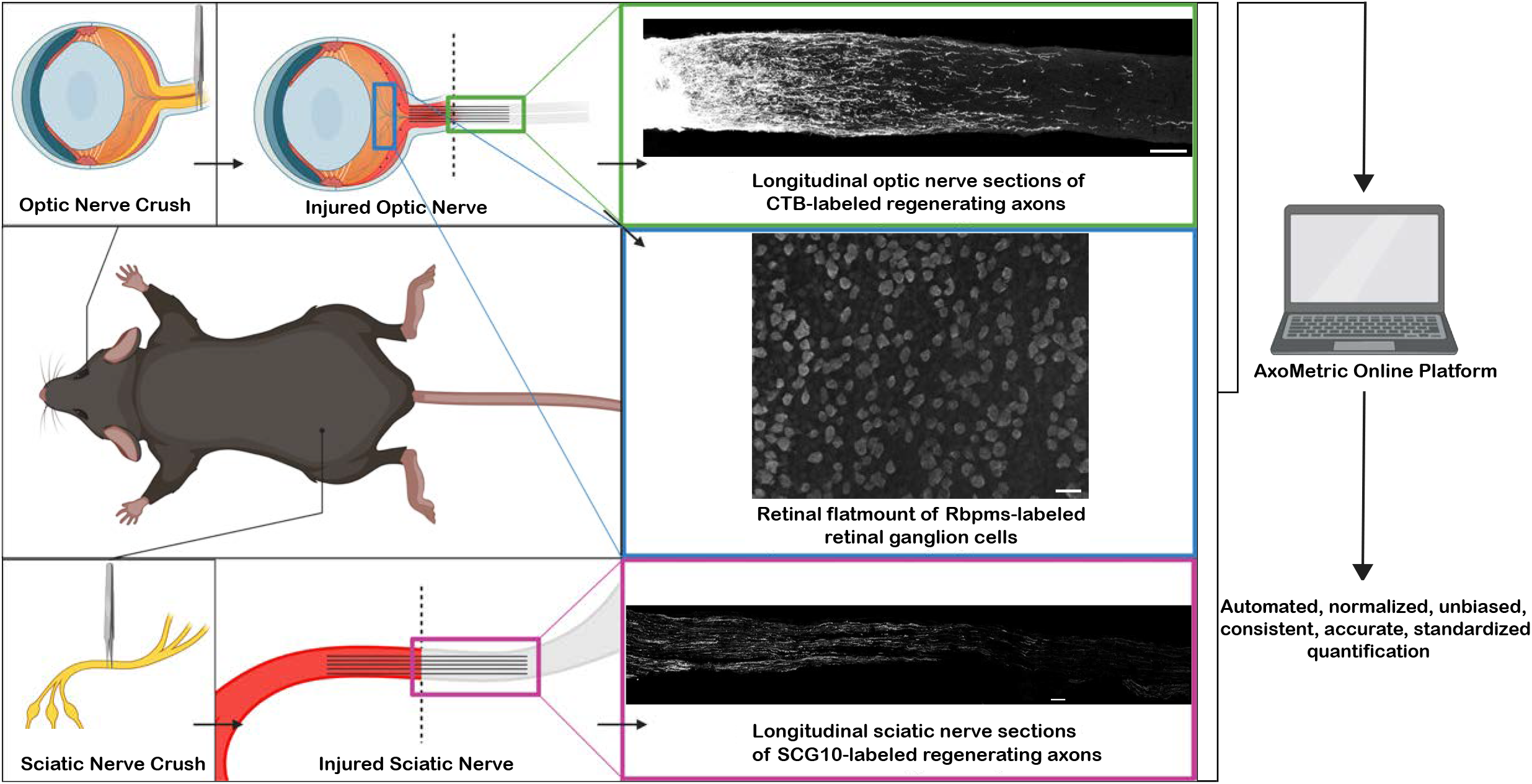

## Introduction

Spontaneous regeneration of severed axons in the adult mammalian central nervous system (CNS) is very limited. One promising method to promote long-distance regeneration of injured retinal ganglion cells (RGC) axons is the activation of specific immune receptors on hematogenous immune cells that accumulate in the posterior chamber of the eye following intraocular injection of crude or highly purified particulate β(1,3)(1,6)glucan (β-glucan).^2,10^ β-glucan is a pathogen-associated molecular pattern (PAMP) ligand recognized by multiple receptors on immune cells, including toll-like receptor 2 (TLR2) and dectin-1 (Clec7a).^2^ In a recent study, we showed that both beneficial and detrimental myeloid cells can positively and negatively influence RGC axon regeneration following intraocular β-glucan injection. Specifically, we found that after retro-orbital optic nerve crush (ONC) injury combined with intraocular β-glucan, microglia protect the inflamed vasculature, reduce leakiness of the blood-retina barrier (BRB), and greatly enhance immune-mediated axon regeneration. In addition, we demonstrated that blocking mature neutrophil trafficking into the eye protects the BRB and greatly enhances immune-mediated axon regeneration.^16^

To identify the immune cells that influence β-glucan–induced retinal ganglion cell (RGC) axon regeneration, we used a combination of global and conditional gene ablation, antibody-mediated blockade of immune cell trafficking, and pharmacological approaches. For unbiased quantification of regenerated axons across different experimental conditions, we developed a new platform, called *AxoMetric*.

*AxoMetric* is a web-based RShiny application that enables interval-based identification and quantification of regenerated axons at multiple distances from the nerve injury site. To account for variability in nerve section width, axon counts are normalized to the nerve diameter at each distance. Compared to conventional manual quantification, *AxoMetric* provides similar results, however, with substantially improved speed and reproducibility. Additionally, because optic nerve crush leads to RGC loss—and some experimental treatments promote the survival of axotomized RGCs—we incorporated functionality for quantifying labeled RGCs in retinal flat mounts. Here, we introduce the core features of *AxoMetric* and provide step-by-step guidance for its use.

## Materials and methods

### Animals and procedures

All procedures involving mice were approved by the Institutional Animal Care and Use Committee at the University of Michigan (Protocol #PRO00011554) and performed in accordance with guidelines developed by the National Institutes of Health. Wild-type (WT) C57BL/6J (RRID:IMSR_JAX:000664) and transgenic mice, *Sarm1/Myd88-5-/-* (RRID:IMSR_JAX:018069), were used. For a detailed description of surgical procedures, euthanasia, and transcardial perfusions, see methods previously described. ^10,16,27^

### Tissue preparation and image collection

Optic nerves and retinas: for visualization of regenerated axons, mice were subjected to intra-ocular injection of cholera toxin subunit B (CTB, Thermo Fischer Scientific #C22843) two days prior to euthanizing. Optic nerves were then embedded in OCT (Tissue-Tek, 4583) and serially sectioned at 14 µm thickness at – 18°C, using a Leica CM 3050S Cryostat. Nerve sections were mounted on Superfrost Plus (Fisherbrand, 12-550-15) microscope slides and stored at -20°C. Nuclei were stained with Hoechst 33343 (Invitrogen #H3570) and sections were mounted in ProLong Gold (Invitrogen, P36930) anti-fade mounting medium. Retinas were dissected and permeabilized in 0.5% TritonX-100 in PBS for 1 hour at room temperature, followed by blocking with 3% donkey serum (EMD Millipore #S30-100ML) and 1% bovine serum albumin (CAS 9048-46-8) for 2 hours at room temperature. Retinas were then incubated with primary antibodies (Rabbit anti-Rbpms Abcam #194213, RRID:AB_2920590, 1:500) rocking overnight at 4 °C. The following day retinas were rinsed 3x with PBS for 30 minutes each and incubated with secondary antibodies (Donkey anti-Rabbit 488 Jackson ImmunoResearch Labs, #711-545-152, RRID:AB_2313584, 1:200) overnight at 4 °C. Optic nerve sections were imaged using the Zeiss Axio Observer Z1 inverted fluorescence microscope fitted with a Zeiss Axiocam 503 mono camera and Zen 3.9 software. The Apotome was enabled for all images at a scan setting of 3 images per z plane. Optic nerve sections were imaged with an HC PL APO 20x/0.75 objective and tiled scan using Zen Blue autofocus software with Hoechst set as the reference channel. Optical sections were taken through the tissue at 1.0 µm z-step size. Retina flat mounts were imaged using Zen Blue autofocus software in a single plan with an HC PL APO 20x/0.75 objective.

Sciatic nerves: sciatic nerves were subjected to nerve crush as previously described^27^ and were harvested at 7 days post-crush, embedded in OCT (Tissue-Tek, 4583), and frozen on dry ice. Sciatic nerves were serially sectioned at 14 µm thickness at –18°C, using Leica CM 3050S Cryostat. Sections were mounted on Superfrost Plus (Fisherbrand, 12-550-15) microscope slides, air-dried overnight, and stored at -20°C. Sections were warmed to room temperature, rinsed twice in 1X PBS for 5 minutes each, and permeabilized in 0.3% triton-x-100/PBS for 10 minutes. For blocking, sections were incubated in 5% donkey serum (EMD Millipore, S30) in 0.1% Triton-x-100/PBS for 1 hour. Sections were incubated with rabbit α-STMN2/SCG10 (Novus Biologicals, NBP1-49461, RRID:AB_10011569, 1:2000) in blocking buffer overnight at 4°C in a humidified chamber. The next day, sections were rinsed three times in 1X PBS, 5 minutes each and were incubated in secondary antibody donkey anti-rabbit 488 (Invitrogen, A-21206, RRID:AB_2535792, 1:1000) in blocking buffer for 2 hours in a humidified chamber. Sections were mounted in Prolong Gold (Invitrogen, P36930) anti-fade mounting medium and imaged. Sciatic nerve sections were imaged using a Zeiss Axio Observer Z1 inverted fluorescence microscope fitted with a Zeiss Axiocam 503 mono camera and Zen 3.9 software. The Apotome was enabled for all images at a scan setting of 3 images per z plane. Sciatic nerve sections were imaged with an HC PL APO 20x/0.75 objective and tiled scan using Zen Blue autofocus software with Hoechst set as the reference channel. Optical sections were taken through the tissue at 1.0 µm z-step size.

### Manual Quantification

Optic nerve images were pre-processed and aligned by the injury site vertically in Microsoft PowerPoint. Four horizontal, equally spaced lines were placed at measurement points relative to the crush site throughout the length of the nerve. Axons were counted within the nerve at each of the intersections between the horizontal lines and in the image (indicating each measurement location for axons relative to the injury site) and were used as manual counts.

Sciatic nerve images were pre-processed and analyzed using FIJI. First the scale is set to the pixel-to-inches ratio of the images. Secondly, a line tool was used to measure the distance where the measurement region of interests (ROIs) will be placed. Next, using the Rectangle tool, 50 µm wide ROIs were set at regular intervals along the nerve length that cover the entire nerve width. The ROI positions were recorded using the ROI manager feature. When all ROIs were placed in the desired locations, the multi-measure tool was used to display the MFI information of each ROI, which were used for manual counts.

Retina flat mount images were pre-processed and analyzed using FIJI. The Multi-point tool was used to count the number of Rbpms-positive cells identified by the user and was used for manual counts.

### Image pre-processing for input

Optic and sciatic nerve images were exported from the Zeiss Version 3.9 software as full-sized TIFF file and then uploaded into FIJI imaging software. Images were straightened and cropped from the start of the injury site to the distal end of the nerve at the same distance for all nerves using the FIJI “Straighten” function. For MFI quantification, users must be careful not to create empty space when cropping images. For quantification, users must make sure the nuclear-stained signal (e.g., Dapi) is not present on the cropped image borders or the code may fail to recognize upper/lower limits. The intensity of signal was adjusted the same across all images to enhance visualization of SCG10-stained or CTB-traced axons, images were scaled to 50% and were then saved as an 8-bit TIFF file (.jpeg files are also compatible with the platform, yet gave slightly different but still comparable quantification results to TIFF files). Retina images were exported from the Zeiss Version 3.9 software as full-sized TIFFs and then uploaded into FIJI imaging software. The intensity of the signal was adjusted the same across all images to enhance visualization of the anti-Rbpms signal, images were scaled to 50% and were then saved as an 8-bit TIFF (.jpeg files are also compatible with the platform, yet gave slightly different but still comparable quantification results to TIFF files).

### User interface

RStudio software (version 4.4.0), the Shiny package (version 1.9.0), and the EBImage package (version 3.20)^18^ were used to generate *AxoMetric* and may be accessed through the following link: https://cdb-rshiny.med.umich.edu/Giger_AxoMetric/. The *AxoMetric* homepage displays descriptions of how the platform works, what tools are available, and where to find these tools to use in the corresponding tabs on the left side of the home page. The first tab “Axon Quantification” has multiple features that can be used for quantification of the “total axons” (CTB-labeled) in an image with a longitudinal optic nerve section; “normalized axons” allows normalization of the total number of axons in an optic nerve image to nerve width; and “multiple files” allows processing and quantification of multiple optic nerve files at once for total axons. The “MFI Quantification” tab has options to quantify the mean fluorescence intensity in nerve images, to normalize the mean fluorescence intensity to nerve width, and to process the quantification of multiple nerve files at once. The “RGC Quantification” tab has the option to quantify the number of retinal ganglion cells (anti-Rbpms staining) in one retina image, or a second option to quantify multiple retina images at once. Users will upload their respective optic nerve, sciatic nerve, and/or retina images to the corresponding tab, submit the image(s) for processing (**Figure 5A**). Different output options of quantified structures are shown (**Figure 5B**). All quantification options provide step-by-step instructions (**Figure 5C**) and results in the export format of Excel or CSV files on their respective pages.

### Software details

#### CTB-based axon quantification

Pre-processed optic nerve images are uploaded (**Figure 1A**) and the CTB signal in each image is dilated to minimize discontinuous gaps of CTB signal for more accurate quantification of axons (**Figure 1B**). Pixel signal thresholding is used to create a mask that consists of a binary array of pixels that are either above or below the optimized CTB-signal threshold (**Figure 1C**). Continuous signals of pixels are grouped into a single object, and objects smaller than the optimized standard size of an axon at 20x are filtered out of the mask (**Figure 1D**). Optimized thresholds of pixel signal strength and object size are intentionally kept non-editable to ensure consistent image analysis across users and to promote reproducibility and standardization across quantification in the field. The final filtered image is then analyzed for continuous signals of pixels that are grouped as objects (traced and color-coded) that may then be used for quantifying of the number of total axons (**Figure 1E**). The total number of axons per section is then converted to the number of axons per nerve according to conventional quantification methods (Σ*a*_*d*_ = π*r*^2^x [average axons/mm]/t)^10^ and by using an estimated nerve radius of 150 micrometers and thickness of 14 micrometers. The result output may then be exported as a(n) CSV/Excel file. The CTB-labeled axon quantification is optimized from longitudinal sections (14 µm) of mouse optic nerves imaged with a 20x objective.

**Figure 1.**
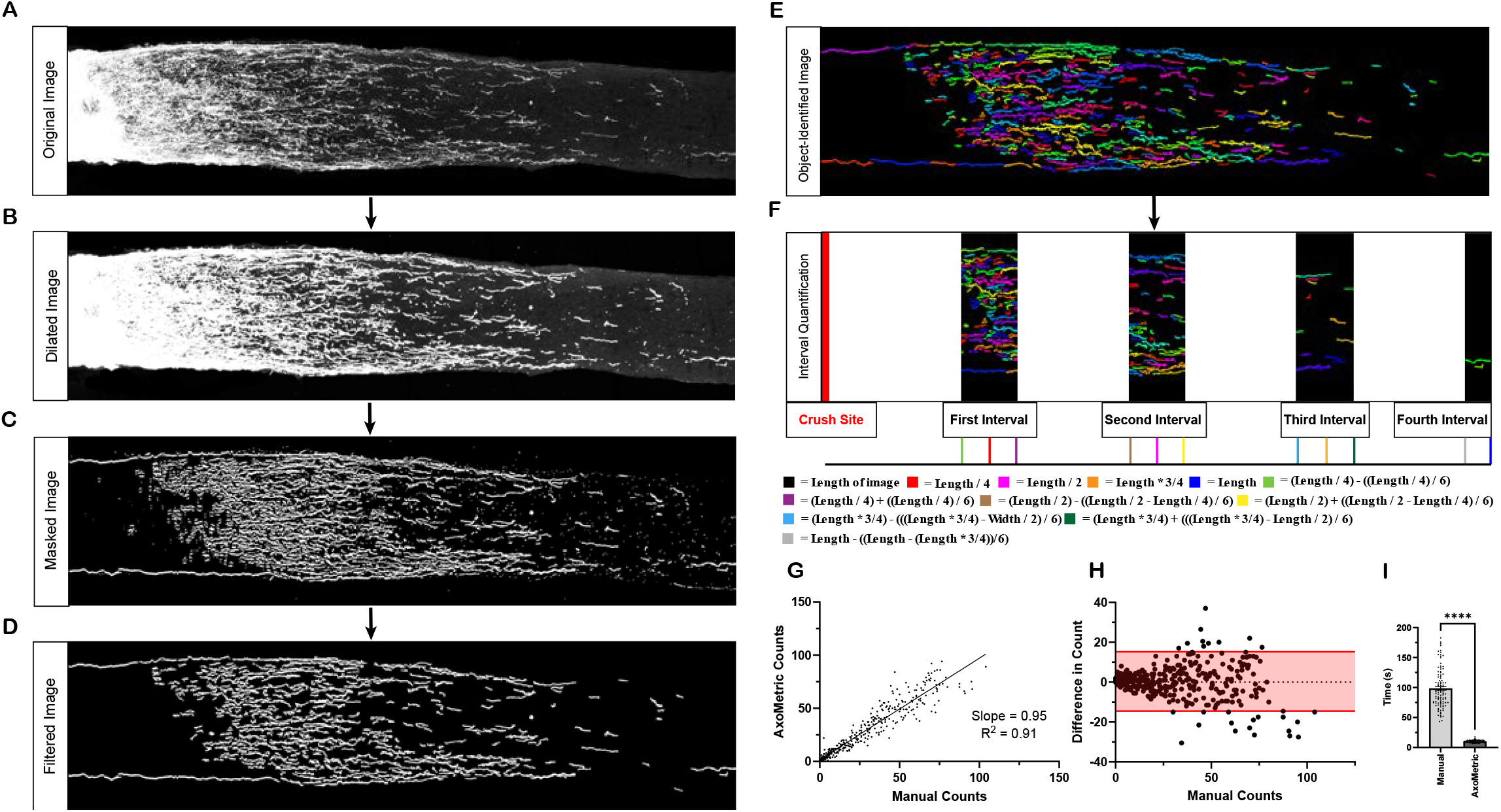
*AxoMetric* interval-based quantification of CTB-labeled regenerating axons in the mouse optic nerve. (A) Original input image of mouse optic nerve labeled with CTB for regenerating axons. (B) Dilated image rendered from the original input image with dilated CTB signal. (C) Masked image created from the dilated image where pixel signal is masked based on cut-off threshold values to identify continuous CTB signal objects. (D) Filtered image generated by selective removal of objects classified below an optimized threshold for size. (E) Colored image indicating individual objects with continuous CTB signal. (F) Interval-based quantification diagram demonstrating the quantification of the total number of axons at four equal intervals relative to the crush site, with their respective distances relative to image length color-coded below. (G) Pearson correlation comparison between manual and automated *AxoMetric* axon counts of mouse optic nerves at four equal interval distances (500, 1,000, 1,500, & 2,000 µm) from the crush site (n = 400), slope = 0.95 & R^2^ = 0.91, ****p ≤ 0.0001. (H) Bland-Altman plot between manual axon counts and the differences between manual and automatic counts of mouse optic nerves at four equal interval distances (500, 1,000, 1,500, & 2,000 µm) from the crush site (n = 400). 95% agreement interval is marked in red, 95% upper limit of agreement = 15.25, 95% lower limit of agreement = -14.49. (I) Comparison between the time needed for manual and automated *AxoMetric* axon counting per optic nerve (n = 100). Quantification of time to count axons per nerve presented as mean +/-SEM, and data was analyzed with Wilcoxon’s t-test, ****p ≤ 0.0001.

#### Mean fluorescence intensity (MFI) quantification

Pre-processed sciatic nerve images (also compatible with optic nerve images) are uploaded (**Figure 2A**), and the fluorescence intensity of each pixel is quantified and averaged across all pixels within their respective interval, where each interval average mean fluorescence is then reported and then may be exported as a(n) CSV/Excel file. The anti-SCG10-labeled axon quantification is optimized from longitudinal sections (14 µm) of mouse sciatic nerves imaged with a 20x objective.

**Figure 2.**
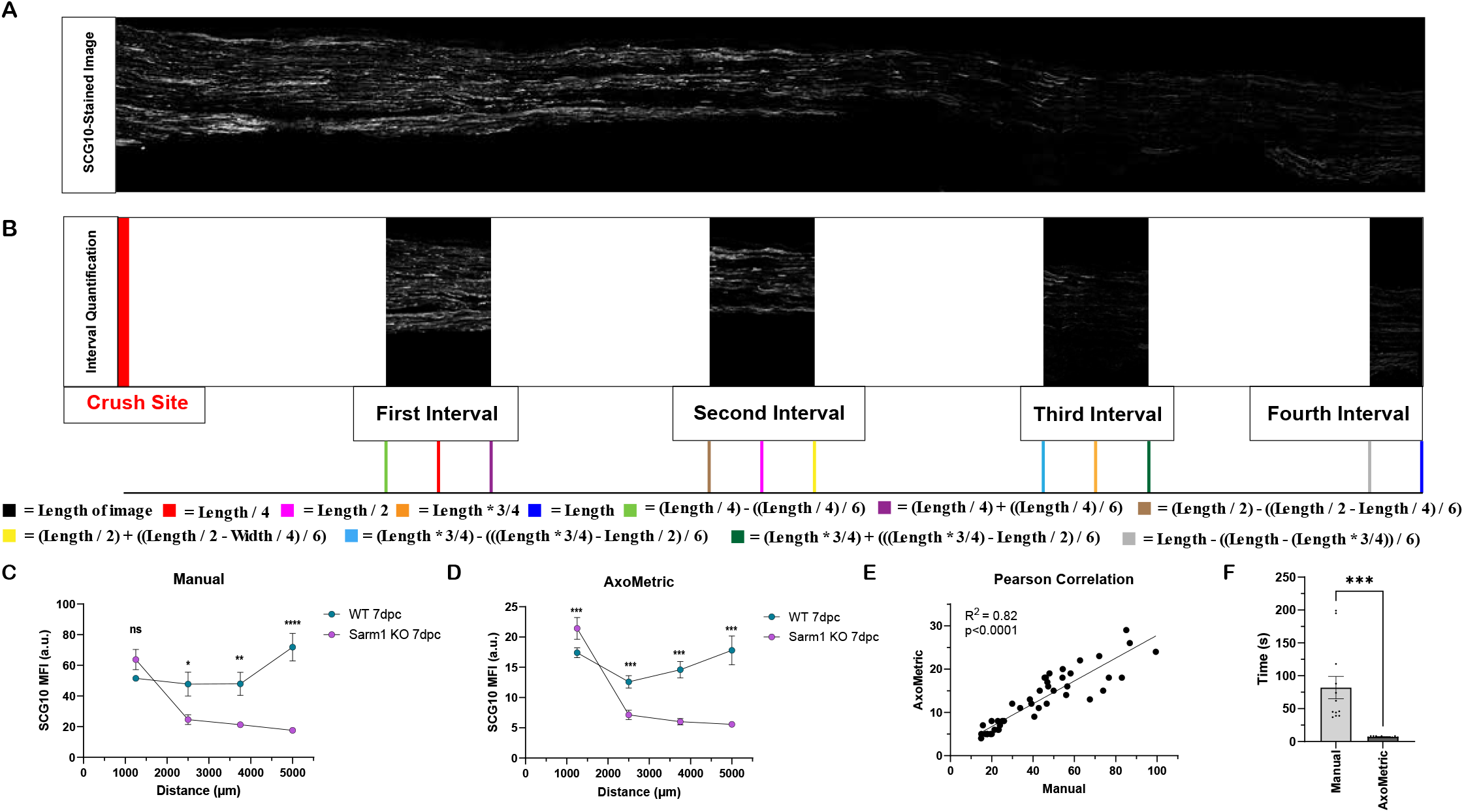
*AxoMetric* interval-based mean fluorescence intensity (MFI) quantification of SCG10-labeled regenerating sensory axons in the mouse sciatic nerve. (A) Original input image of mouse sciatic nerve labeled with SCG10 for regenerating sensory axons. (B) Interval-based quantification diagram demonstrating the quantification of the mean MFI at four equal intervals relative to the crush site, with their respective distances relative to image length color-coded below. (C) Manual quantification of mean SCG10 MFI utilizing FIJI software of wild-type (WT) and Sarm1 Knockout (Sarm1 KO) 7 days post-sciatic nerve crush (n = 5 & 7 respectively). MFI presented as mean +/-SEM. One-way ANOVA followed by Tukey’s multiple comparisons test. *p ≤ 0.05, **p ≤ 0.01, ****p ≤ 0.0001. (D) Automated quantification of SCG10 MFI utilizing *AxoMetric* software of WT and Sarm1 KO 7 days post-sciatic nerve crush (n = 5 & 7, respectively). Quantification of MFI presented as mean +/-SEM. One-way ANOVA followed by Tukey’s multiple comparisons test. ***p ≤ 0.001. (E) Pearson correlation comparison between manual and *AxoMetric* mean MFI quantification of mouse sciatic nerves at four intervals relative to the crush site, R^2^ = 0.82, ****p ≤ 0.0001. (F) Comparison between the time needed for manual and *AxoMetric*’s automated mean fluorescence intensity quantification per sciatic nerve (n = 12). Time to quantify mean fluorescence intensity per nerve presented as mean +/-SEM and data was analyzed with Wilcoxon’s t-test, ****p ≤ 0.0001.

**Figure 3.**
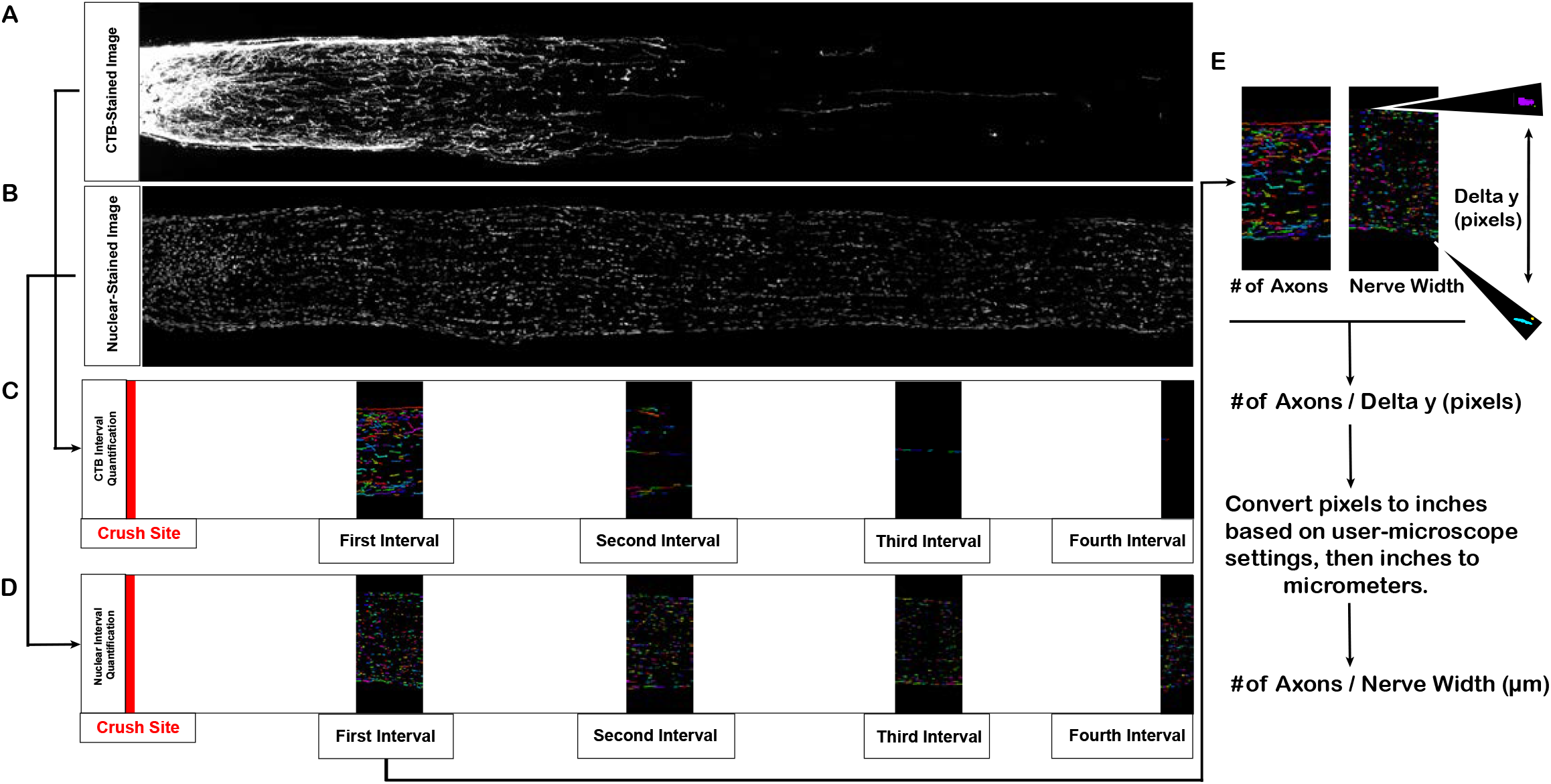
*AxoMetric* interval-based quantification and normalization of CTB-labeled regenerating axons to nerve width. (A) Mouse optic nerve longitudinal section stained with CTB for regenerating axons. (B) Mouse optic nerve longitudinal section stained with a nuclear stain (Hoechst) to identify nuclei. (C) Interval-based quantification of CTB-stained regenerating axons at four equal intervals relative to the crush site. (D) Interval-based quantification of Hoechst-stained nuclei at four equal intervals relative to the crush site. (F) Diagram of the calculation and normalization of the number of axons per nerve width.

#### Anti-Rbpms-based retinal ganglion cell quantification

Pre-processed retina images are uploaded, and the pixel signal of Rbpms staining is thresholded to create a mask that consists of a binary array of pixels above an optimized pixel signal for surviving retinal ganglion cells. Continuous signals of pixels are grouped into objects in the final image and is analyzed for continuous signals of pixels that are grouped as objects (color-coded) that may then be used for quantification of retinal ganglion cells (**Figure 4A**) and then may be exported as a(n) CSV/Excel file. The anti-Rbpms-labeled retinal ganglion cell quantification is optimized for flat-mounts of mouse retinas imaged with a 20x objective.

**Figure 4.**
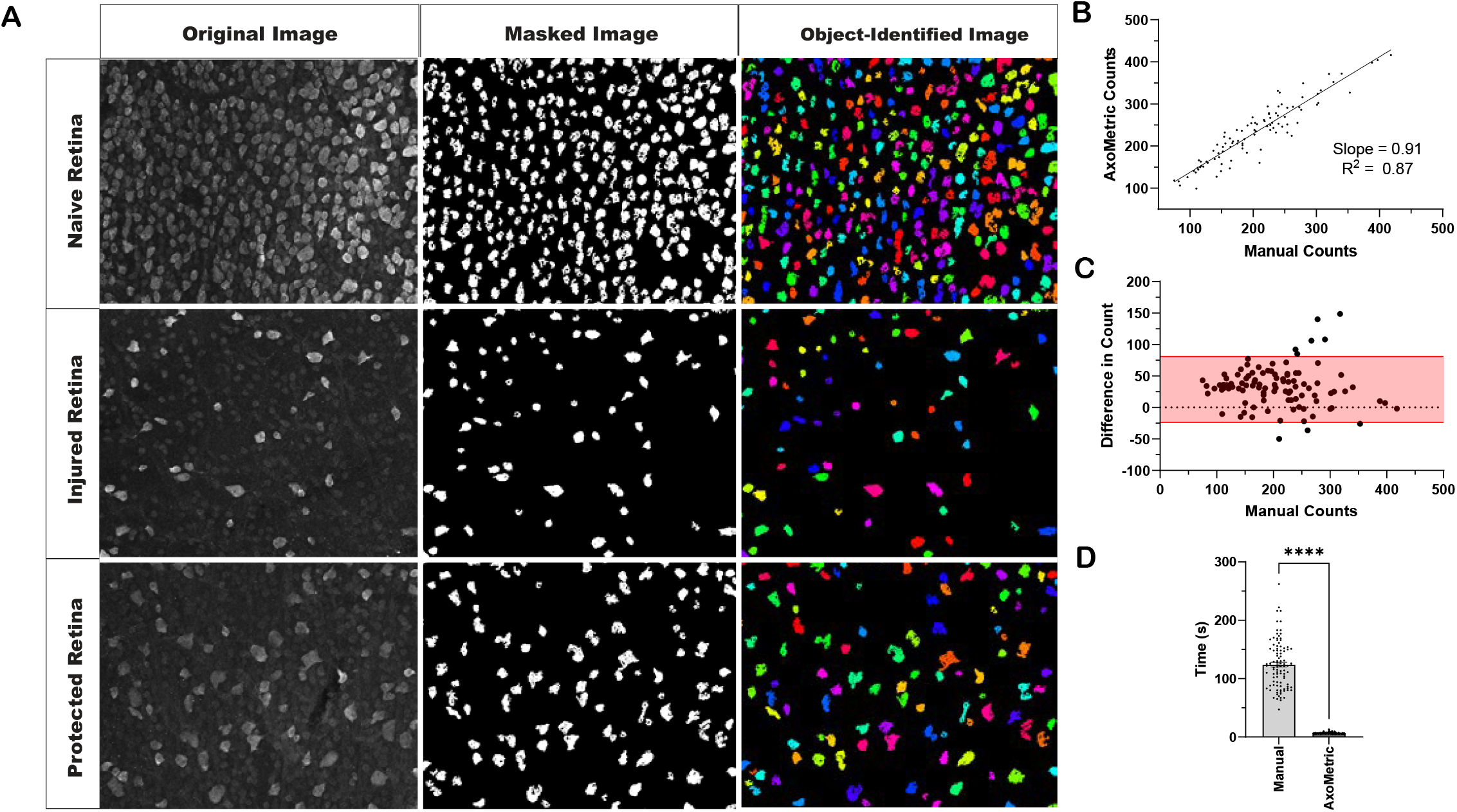
*AxoMetric* quantification of Rbpms-labeled retinal ganglion cells in the mouse retina. (A) Naïve, injured (optic nerve crush 14 days post crush), and protected (conditioned lens injured and optic nerve crush 14 days post crush) retinas labeled with Rbpms for quantification of retinal ganglion cells. Original input images, masked images based on pixel signal intensity threshold, and object-identified images of objects with continuous Rbpms signal. (B) Pearson correlation comparison between manual and *AxoMetric*’s automated retinal ganglion cell counts mouse naïve, injured, and protected retina images (n = 100). Slope = 0.91 & R^2^ = 0.87, ****p ≤ 0.0001. (C) Bland-Altman plot between manual retinal ganglion cell counts and the differences between manual and *AxoMetric*’s automatic counts of mouse retinas (n = 100). 95% upper limit of agreement = 80.98, 95% lower limit of agreement = -23.65. (D) Comparison between the time needed for manual and *AxoMetric*’s automated retinal ganglion cell counting per retina image (n = 100). Quantification of time to count retinal ganglion cells per retina image presented as mean +/-SEM. Wilcoxon’s t-test, ****p ≤ 0.0001.

**Figure 5.**
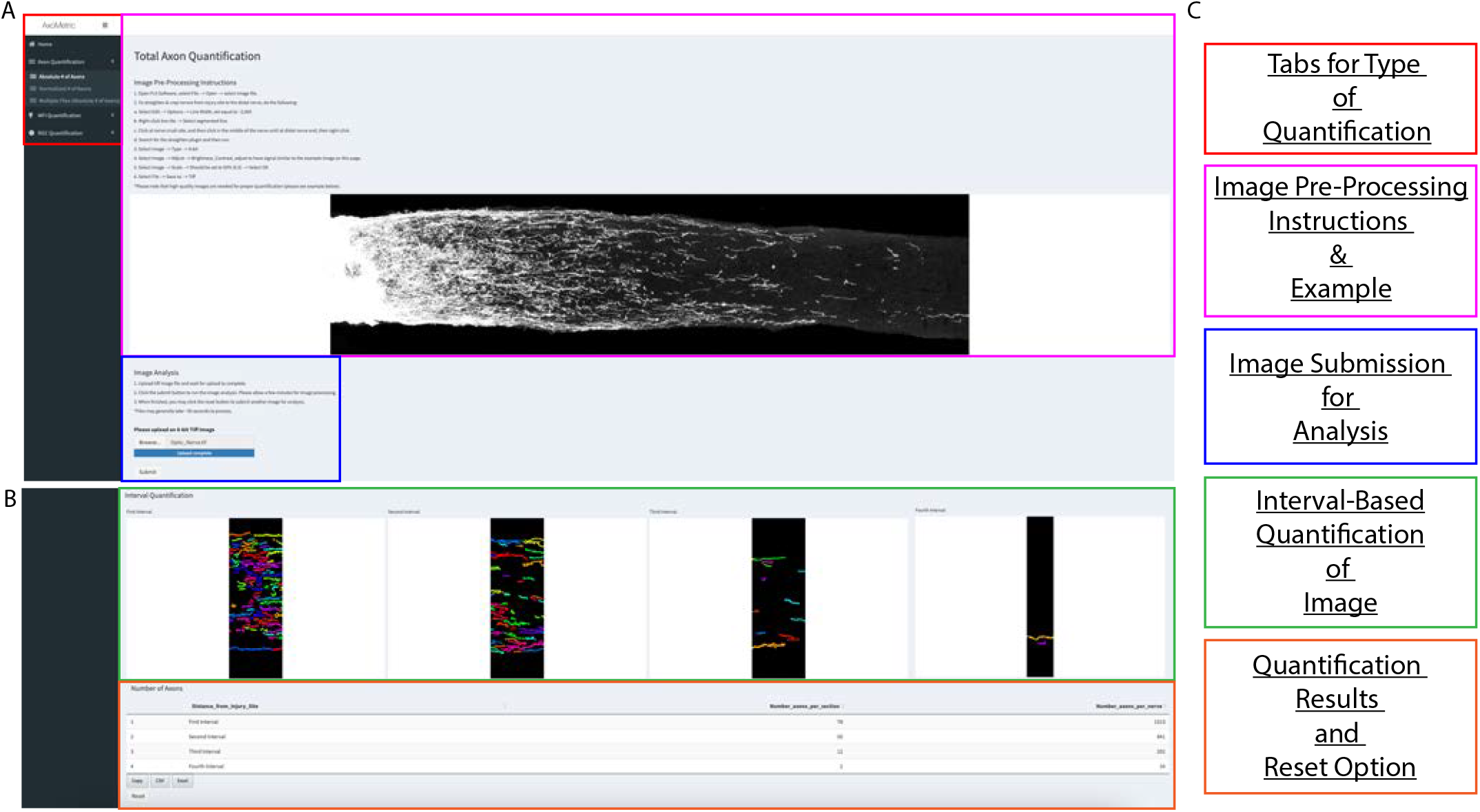
*AxoMetric* user interface example for automatic interval-based quantification of a mouse optic nerve. *(A) AxoMetric* user interface for selection of the type of quantification, image pre-processing instructions and examples, and image submission for analysis. *(B) AxoMetric* user interface for interval-based quantification of the image, quantification results, and reset options. (C) General workflow of the *AxoMetric* user interface labeled by section.

#### Interval-based quantification

Optic and sciatic nerve images are pre-processed to be cropped from the injury site to a distal site that is equal in distance for all images being quantified in the same experiment. The code uses the image length to determine interval distances. For example, researchers may use mouse optic nerves cropped to 2.0 mm distal to the injury site and quantify intervals at 0.5, 1.0, 1.5, and 2.0 mm respectively, or a researcher may crop 4.0 mm distal to the injury site for all nerves, and quantify intervals at 1.0, 2.0, 3.0, and 4.0 mm respectively. Images are either quantified according to the number of axons or mean fluorescence intensity descriptions above, and then four equal intervals are created in the image relative to the crush site and the length of the image – which may be adjusted to fit four equal quantification intervals at any length (**Figure 1F** and **Figure 2B**). The center of the first interval is the length of the image divided by four, the center of the second interval is the length of the image divided by two, the center of the third interval is the length of the image multiplied by ¾, and the center of the last interval is the end of the image. The optimized upper and lower length limits of each interval with respect to the center of the interval are 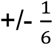 of the interval length on either side of the center. Once the four equal intervals are calculated, either the number of objects identified from CTB for axons, or the mean fluorescence intensity, is quantified for each respective interval.

#### Normalization to nerve width

Optic and sciatic nerve images are pre-processed for stains of interest (e.g., CTB or SCG10, **Figure 3A**) and require inclusion of nuclear staining to determine nerve width (Hoechst, **Figure 3B**). Both the stain of interest and nuclear-stained images of the nerves are uploaded, and the stain of interest image is processed following the interval-based quantification of either CTB-labeled axons or mean fluorescence intensity (**Figure 3C**). The nuclear-stained image is then processed in the same way as the stain of interest, except the code is optimized to identify nuclei-related objects at 20x (**Figure 3D**). At each interval, the maximum and minimum nuclei (analyzed by relative pixel y-positions in the image) are selected and the distance between the nuclei is measured in pixels (Delta y, **Figure 3E**). The user is prompted to provide the image’s pixel height (in inches), found in the image properties tab of FIJI, and then the code may convert pixels to inches and then inches to micrometers. After calculating the nerve width (i.e., diameter) of the nerve at each interval in micrometers, this may then be used with the respective axon or mean fluorescence intensity measurements (described previously, except substituting in the calculated nerve radius by dividing the nerve width in half) to determine the number of axons or MFI per nerve width in an interval-based manner, correcting for anatomical variations.

### Statistical Analysis

Data are presented as mean ± SEM. Statistical analysis was performed in GraphPad Prism (v10) using Wilcoxon’s signed-rank *t* test, Pearson correlation, and Bland-Altman analysis, as indicated in the figure legends. A *p value<0.05 was considered significant. **p<0.01, ***p<0.001, ****p<0.0001. Data acquisition and analysis were carried out by individuals blinded to experimental groups.

## Results

*AxoMetric* is a novel, user-friendly, open-source platform that enables rapid and unbiased quantification of labeled structures in tissue sections. Designed with a focus on neural tissues, the platform supports multiple applications, including quantification of regenerated axons in the optic nerve via individual axon tracing, bundled axon density in the sciatic nerve using mean fluorescence intensity (MFI), and retinal ganglion cell density in flat mounts. By eliminating the need for time-consuming manual quantification, *AxoMetric* enhances both efficiency and reproducibility. The software is compatible with both PC and Macintosh desktop or laptop systems and includes features such as normalization to nerve width, which allows for more accurate and standardized measurements. By streamlining and standardizing regeneration quantification, it is our hope that *AxoMetric* enables assessment of regeneration efficacy, across different treatments strategies and different laboratories.

### AxoMetric Accurately Recapitulates Manual Axon Counts in the Optic Nerve with Significantly Increased Speed

To determine the accuracy and efficiency of *AxoMetric’s* software to automatically count regenerated axons in the optic nerve at four intervals distal to the crush site, 100 mouse optic nerves were counted with *AxoMetric* and then compared to averaged manual counts of the same optic nerve sections (counts were averaged between two nerve sections of the same nerve) by two independent investigators. A Pearson correlation analysis was performed between *AxoMetric’s* axon counts and the two manual counts (R^2^ = 0.89 and a slope of 0.74 – investigator 1, R^2^ = 0.87 and a slope of 1.25 – investigator 2), yielding an averaged R^2^ = 0.91 and a slope of 0.95 (**Figure 1G**). A Bland-Altman comparison between manual and *Axo*Metric’s axon counts demonstrated that 94% of automated counts from *AxoMetric* are within the 95% limit of agreement with manual counts (**Figure 1H**). *Axo*Metric’s axon counts were more accurate with lower manual counts, and more dispersed with higher manual counts, likely due to the blending of CTB signal with large numbers of axons near the crush site in the first interval of quantification. The median time for *AxoMetrics* automated counting was 10.00 seconds to count axons per optic nerve section compared to 98.46 seconds for manual counting (**Figure 1I**), significantly reducing the time to count axons per optic nerve, excluding image pre-processing time (p ≤ 0.0001, Wilcoxon t-test). These results suggest that *Axo*Metric can quantify CTB-labeled regenerating axons in mouse optic nerves accurately and more efficiently than manual axon counting.

### Application of AxoMetric for Fluorescence Intensity Quantification of Regenerated Axons in the Sciatic Nerve

Regenerating axons in the injured mouse sciatic nerve form tightly fasciculated bundles, also called Bands of Bungner, making individual axon counting impractical—unlike in the optic nerve. To address this, we used *AxoMetric* to quantify mean fluorescence intensity (MFI) of anti-SCG10-labeled regenerating sensory axons in sciatic nerve sections harvested 7 days post-crush injury. Quantification was performed at four intervals distal to the crush site and compared to manual MFI measurements using regions of interest in FIJI (**Figure 2C–D**). We analyzed sections from wild-type (WT, *n* = 5) and *Sarm1* knockout mice (*Sarm1*^*-/-*,^ *n* = 7) in which Wallerian degeneration is delayed and causes a regeneration phenotype.^7^ While the magnitude of mean fluorescence intensity absolute unit measurements was different between manual and *AxoMetric* counts, axon regeneration was significantly decreased in the *Sarm1*^*-/-*^ distal nerve. This shows that *AxoMetric* accurately captured the previously reported regeneration defects in the Sarm1^*-/-*^ mice.^23^ A Pearson correlation analysis was performed between *AxoMetric’s* automated and manual MFI quantification, yielding an R^2^ = 0.82 (**Figure 2E**). The average post-image processing time for *AxoMetric’s* automated quantification was 7.170 seconds compared to 82.17 seconds for manual quantification (**Figure 2F**), significantly reducing the time to quantify sciatic nerve regeneration, excluding image pre-processing time (p ≤ 0.0001, Wilcoxon t-test). These results show that *AxoMetric* can, in an unbiased and consistent fashion, quantify anti-SCG10 labeled regenerating sensory axon bundles in mouse sciatic nerves accurately and faster than manual quantification.

### AxoMetric Enables Efficient and Accurate Counting of Retinal Ganglion Cells

We developed an automated counting module within *AxoMetric* for quantifying retinal ganglion cell (RGC) nuclei labeled with anti-RBPMS, similar in principle to existing FIJI-based methods. To assess its accuracy, we analyzed 100 retinal flat mounts from naïve and optic nerve–injured mice, with and without neuroprotective treatments (e.g., conditioning lens injury)^10^. *AxoMetric* was used, and the results compared to the averaged manual counts performed by two independent investigators, blinded by experimental treatment. Pearson correlation analysis between *AxoMetric* and both manual counts (R^2^= 0.89 and slope = 0.99 – investigator 1, R^2^= 0.73 and slope = 0.74 – investigator 2) yielded an averaged correlation of R^2^= 0.87, slope = 0.91 (**Figure 4B**). Bland-Altman analysis showed that 92% of *AxoMetric* counts fell within the 95% limits of agreement with manual quantification (**Figure 4C**). The median time required for automated RGC counting was 6.90 seconds per retina, compared to 124.1 seconds for manual analysis (**Figure 4D**), representing a significant reduction in processing time, excluding image pre-processing time (p ≤ 0.0001, Wilcoxon test). These findings demonstrate that *AxoMetric* accurately and efficiently quantifies anti-RBPMS–labeled RGCs in retinal flat mounts, providing a reliable alternative to manual counting.

## Discussion

Previous software platforms have been developed to quantify protected and regenerated neurons in the central and peripheral nervous systems. Early tools developed for RGC and DRG neuron axon quantification (e.g., AxonMaster, AxonJ, Axonet, AxonDeep, AxoNet2.0, AxoDetect) utilized axial rather than longitudinal nerve sections, limiting the ability to accurately measure axonal distance from the injury site.^1, 4, 6, 8, 12, 21, 22, 25^ Other methods, including NeuronJ, RGC-NET, AxonTracer, and AxonDeepSeg rely on tracing techniques to quantify neurites or axons in *in vitro* culture systems^13, 20^ or in injured spinal cord tissue.^15, 17, 24^ While useful, these approaches do not support linear, interval-based quantification of axons from a defined injury site as implemented in *AxoMetric*.

AxonQuantifier, a plugin-based tool, performs gray-value quantification in an interval-based manner from the injury site.^19^ However, it is only semi-automated, requires user input for defining intervals, and does not support quantification of total axon counts or mean fluorescence intensity in a fully automated, interval-based format. Moreover, it lacks the ability to normalize axonal measurements to nerve width using nuclear staining, a key feature of *AxoMetric. AxoMetric*’s quantification of immunofluorescently labeled RGC nuclei (anti-RBPMS) is comparable to manual quantification described previously.^3, 5, 9, 11, 14, 26^

In sum, *AxoMetric* is a free, publicly accessible Shiny-based platform designed for automated, accurate, and efficient interval-based quantification of regenerating axons, mean fluorescence intensity of axon fascicles, and labeled RGCs in an unbiased manner. It provides researchers with an integrated tool for analyzing stained images of optic or sciatic nerve sections, normalizing to nerve width, and simultaneously quantifying multiple image types within a unified interface. *AxoMetric* is a novel platform suitable for both central nervous system (retina/optic nerve) and peripheral nervous system (sciatic nerve) models, offering standardized, reproducible, and scalable quantification of neuroprotection and axon regeneration.

### Limitations

*AxoMetric* provides researchers with a valuable tool for automating and standardizing the quantification of regenerating axons and labeled cell somas. However, certain limitations should be acknowledged. In CTB-labeled optic nerve images, discontinuous or punctate CTB signal may be interpreted as multiple axons rather than a single continuous structure, potentially leading to overestimation in automated counts. To minimize this overestimation, image signal is dilated in our platform to reduce the amount of discontinuous or punctate CTB signal. Variability was high proximal to the injury site, where axonal density is high, and signal overlap is common. Additionally, the current version limits interval-based quantification to four equally spaced intervals distal to the injury site, whereas allowing users to customize the number and spacing of intervals could enhance flexibility. In cases where more intervals are needed for quantification, the user may crop images in two halves for separate analysis (e.g., four intervals into eight intervals for a single nerve divided into two halves). Another improved feature would be to enable batch quantification for the nerve width normalization quantification method, and to recommend processing more than ten nerve images at a time, however, this is currently limited by server capacity. The current capacity only allows for one user to process images at a time, otherwise, other users will receive an error message until the original user if finished with their analysis. Future directions include adapting the platform for use with other tissues and species (e.g., spinal cord and rat), integrating artificial intelligence to further improve the accuracy of axon/cell soma quantification, and increasing the server’s bandwidth to accommodate more than one user over time.

## Acknowledgments

We thank Biorender.com for the figures used in this publication (BioRender. Finneran, M. (2025) https://BioRender.com/3mtflv4). We also thank the members of the Giger laboratory and Dr. Larry Benowitz for their assistance in reviewing the manuscript. This work was supported by the National Institutes of Health (NIH) through the Ruth L. Kirschstein National Research Service Award (1F31EY036280-01 to M.C.F.), the Dr. Miriam and Sheldon G. Adelson Medical Research Foundation, and the Stein Innovation Award for Research to Prevent Blindness. The content is solely the responsibility of the authors and does not necessarily represent the official views of the National Eye Institute or other funding agencies. This research was, in part, also funded by the Advanced Research Projects Agency for Health (ARPA-H, R.J.G.). The views and conclusions contained in this document are those of the authors and should not be interpreted as representing the official policies, either expressed or implied, of the United States Government. The funding sources had no role in the study design, data collection and analysis, interpretation of results, manuscript preparation, or the decision to submit the article for publication.

## Authorship Contributions

**Matthew C. Finneran:** Conceptualization, funding acquisition, software development, methodology, investigation, data curation, formal analysis, project administration, supervision, writing - original draft, writing – review & editing. **Tara Rhamani:** Investigation, data curation, formal analysis. **Ismaël Valentin Salioski:** Conceptualization, software development, methodology. **Ligia B. Schmitd:** Investigation, data curation, formal analysis. **Ryan Passino:** Conceptualization, methodology, data curation. **Craig N. Johnson:** Software development, methodology. **Roman J. Giger:** Funding acquisition, project administration, supervision, writing – reviewing & editing.

## Data and Code Availability

Software code available through Github: https://github.com/GigerLab/AxoMetric

Website: https://cdb-rshiny.med.umich.edu/Giger_AxoMetric/

Any additional information required is available from the lead contact upon request.

